# Allele specific expression analysis identifies regulatory variation associated with stress-related genes in the Mexican highland maize landrace Palomero Toluqueño

**DOI:** 10.1101/152397

**Authors:** M. Rocío Aguilar-Rangel, Ricardo A. Chávez Montes, Eric Gonzalez-Segovia, Jeffrey Ross-Ibarra, June K. Simpson, Ruairidh J. H. Sawers

**Author notes:** These authors contributed equally to this work (M. Rocío Aguilar-Rangel, Ricardo A. Chávez Montes). Corresponding Author: Ruairidh Sawers.

## Abstract

**Background:** Gene regulatory variation has been proposed to play an important role in the adaptation of plants to environmental stress. In the central highlands of Mexico, farmer selection has generated a unique group of maize landraces adapted the challenges of the highland niche. In this study, gene expression in Mexican highland maize and a reference maize breeding line were compared to identify evidence of regulatory variation in stress-related genes. It was hypothesised that local adaptation in Mexican highland maize would be associated with a transcriptional signature observable even under benign conditions.

**Methods:** Allele specific expression analysis was performed using the seedling-leaf transcriptome of an F1 individual generated from the cross between the highland adapted Mexican landrace Palomero Toluqueño and the reference line B73, grown under benign conditions. Results were compared with a published dataset describing the transcriptional response of B73 seedlings to cold, heat, salt and UV treatments.

**Results:** A total of 2386 genes were identified to show allele specific expression. Of these, 277 showed an expression difference between Palomero Toluqueño and B73 alleles that mirrored the response of B73 cold, heat, salt and/or UV treatments, and, as such, were considered to display a constitutive stress response. Constitutive stress response candidates included genes associated with plant hormone signaling and a number of transcription factors. Construction of a gene co-expression network revealed further signaling and stress-related genes to be among the potential targets of the transcription factors candidates.

**Discussion:** Constitutive activation of responses may represent the best strategy when stresses are severe but predictable. Expression differences observed here between PT and B73 alleles indicate the presence of *cis*-acting regulatory variation linked to stress-related genes in PT. Considered alongside gene annotation and population data, allele specific expression analysis of plants grown under benign conditions provides an attractive strategy to identify functional variation potentially linked to local adaptation.

## INTRODUCTION

Extensive study across different plant species has identified a range of transcriptional responses to abiotic stresses. Although basic responses are typically conserved, variation in the regulation of stress-responsive genes has been observed among individuals and varieties, potentially playing an important role in adaptation to stressful environments (Hannah et al., 2006); (Swanson-Wagner et al., 2012); (Rengel et al., 2012); (Lasky et al., 2014). From an agronomic perspective, biotechnological approaches to enhance crop stress tolerance to abiotic stress often aim to manipulate gene expression rather than engineer protein sequences (*e.g.* (Kamthan et al., 2016)). Similarly, efforts to identify suitable material for breeding towards similar goals have drawn on natural *cis*-acting regulatory variation acting on stress-responsive gene expression (*e.g.* (Mao et al., 2015)). As such efforts are intensified in the face of mounting concern regarding the impact of climate change on crop productivity, there is ever greater interest in the genetic basis of variation in stress-responses (Des Marais, Hernandez & Juenger, 2013).

Crop landrace varieties represent an invaluable genetic resource. Collectively, the range of environments exploited by landraces typically exceeds that of improved varieties, and many landraces are adapted to conditions that would be considered stressful in conventional agriculture (Ruiz Corral et al., 2008) (Romero Navarro et al., 2017). Nonetheless, although landraces represent a compelling source for enhancing abiotic stress tolerance in breeding programs, the task of identifying useful genetic variants and transferring them to breeding material is far from trivial (Sood et al., 2014). In addition to the complication of working with often heterogenous landrace germplasm, reproducing stress conditions for evaluation is costly and difficult. Furthermore, stress is not well reflected by a single experimental treatment, but rather represents a continuous environmental range defined by interacting variables acting over the lifetime of the plant. Large-scale phenomics efforts are an attempt to implement the factorial designs required to capture such complexity (Houle, Govindaraju & Omholt, 2010); (Furbank & Tester, 2011), but they require a substantial investment in infrastructure that may not be feasible in many research contexts. One possible alternative is to leave aside the difficulties of managing the environment and to look for signatures of an enhanced stress response that are hardwired in locally adapted material and evident under benign conditions.

Stress responses are considered to be an adaptation to an unpredictable, changing environment. When conditions are adverse, but predictable, however, theory suggests that the plastic response may be replaced by constitutive activation, a process referred to as canalization (Waddington, 1942); (Levins); (von Heckel, Stephan & Hutter, 2016). One potential advantage of canalization is to avoid the delay between stimulus and response inherent in plasticity. In cultivated systems, non-adapted varieties can benefit from mild priming stress treatments that activate protective mechanisms and prepare the plants for future more severe environmental challenges (Hilker et al., 2016). In practice, however, the first exposure to a stress may be severe, placing the unprepared organism at risk. Under strong yet predictable environmental stress, constitutive activation may represent the best strategy. On this basis, the transcriptome may reflect local adaptation even under benign conditions, presenting an opportunity to identify genetic variation related to enhanced stress tolerance without the complication of managing the stress environment.

Comparative transcriptome analysis of stress tolerant and non-tolerant varieties provides a powerful approach to identify the molecular mechanisms underlying tolerance variation (*e.g.* (Hayano-Kanashiro et al., 2009); (von Heckel, Stephan & Hutter, 2016). The number of differentially accumulating transcripts, however, may be large, and the data reflect both *cis*-acting and *trans*-acting regulatory variation. Critically, *per se* comparison of varieties has little power to characterize the genetic architecture of stress tolerance or to identify causative genetic variation. In addition, when material is diverse, phenological differences can make it difficult to devise an appropriate sampling strategy. With the development of sequencing based methods to study the transcriptome, it is possible to make use of natural sequence variation to quantify allele specific expression (ASE) in F_1_ hybrid individuals generated from the cross of two different lines of interest (Springer & Stupar, 2007b); (Springer & Stupar, 2007a); (Zhang & Borevitz, 2009); (Lemmon et al., 2014). Characterization of ASE in F_1_ material avoids the problems of comparing parents that may be very different in growth and development by evaluating both alleles within the same cellular environment, directly revealing *cis*-acting genetic variation for transcript accumulation (Springer & Stupar, 2007b); (Lemmon et al., 2014); (Waters et al., 2017).

In this study, a transcriptome dataset was examined for evidence of *cis*-regulatory variation linked to stress-associated genes in Palomero Toluqueño (PT), a maize landrace adapted to the highlands of Central Mexico (Prasanna, 2012); (Perales & Golicher, 2014). The Mexican highland environment exposes maize plants to a number of abiotic stresses: bringing plants to maturity under low-temperatures necessitates planting early in the year, exposing seedlings to late frosts and water deficit before onset of the annual rains; throughout the growing season, low-temperature, high-levels of UV radiation and hail storms pose further challenges (Eagles & Lothrop, 1994); (Lafitte & Edmeades, 1997); (Jiang et al., 1999); (Mercer, Martínez-Vásquez & Perales, 2008); (Ruiz Corral et al., 2008). To identify evidence of regulatory variation that might underlie adaptation to these conditions, an F1 was generated between PT and the midwest-adapted maize reference line B73, and the leaf transcriptome analyzed under benign greenhouse conditions to detect ASE. Results of the analysis were compared with a published study in which B73 seedlings were exposed to cold, heat, salt and UV stress treatments (Makarevitch et al., 2015). A total of 277 genes were identified showing a pattern of ASE under benign conditions that mirrored the response of the same gene under stress in B73, hereafter referred to as constitutively stress responsive (CR). The CR candidate set included transcription factors and genes associated with plant hormone signalling, a number of which are discussed in more detail and presented as candidates for future functional analysis.

## MATERIALS AND METHODS

### Plant material, RNA preparation, and sequencing

Seed of the Mexican highland landrace Palomero Toluqueño accession Mexi5 was obtained from the International Maize and Wheat Improvement Center (CIMMYT; stock GID 244857). The original collection was made near to the city of Toluca, in Mexico state (19.286184 N, -99.570871 W), at an elevation of 2597 masl. An F1 hybrid stock was generated from the cross between the inbred line B73 and PT grown under greenhouse conditions and total RNA was extracted from a single, 14 days-old seedling using the Qiagen RNeasy Plant Mini Kit (cat ID 74904) according to the manufacturer’s protocol. RNA integrity was assessed by spectrophotometry and agarose gel electrophoresis. Library preparation was performed using the Illumina protocol as outlined in the TruSeq RNA Sample Preparation Guide (15008136 A, November 2010) and paired-end sequencing was carried out on the Illumina HiSeq 2000 platform. Raw data is available in the NCBI (www.ncbi.nlm.nih.gov) Sequence Read Archive under accession SRP011579.

### Allele Specific Expression (ASE) analysis

Allele specific expression (ASE) analysis was based on the method of Lemmon and collaborators (Lemmon et al., 2014) and the detailed pipeline is presented as Supplementary Data 1 [pipeline]. A set of 39475 B73 transcripts was generated by selecting the longest predicted transcript for each gene annotated in the AGPv3.22 B73 reference genome (ftp://ftp.ensemblgenomes.org/pub/release-22/). Six transcripts whose sequences consisted of only, or mostly, undefined (N) bases were removed (GRMZM2G031216_T01, GRMZM2G179334_T01, GRMZM2G307432_T01, GRMZM2G316264_T01, GRMZM2G406088_T01 and GRMZM2G700875_T01), resulting in a set of 39469 sequences. A total of 151,168,196 paired-end reads from the B73xPT F1 transcriptome were trimmed using Trimmomatic (Bolger, Lohse & Usadel, 2014) and aligned using bwa mem (Li, 2013) to the set of B73 transcripts. The resulting alignment was processed using samtools, bcftools and vcfutils (Li et al., 2009); (Li, 2011a); (Li, 2011a, b) to identify polymorphisms. We then created a set of PT pseudo-transcripts by substituting the identified sequence variants into the B73 reference transcripts. A single fasta file was created that contained two sequences per locus, one B73 transcript and one PT pseudo-transcript, and B73xPT F1 reads were re-aligned to this F_1_ pseudo-reference using bowtie2 (Langmead & Salzberg, 2012) with eXpress (Roberts et al., 2011); (Roberts & Pachter, 2013) recommended parameters. The number of reads per B73 and/or PT transcript was then quantified using eXpress. A total of 9256 transcripts were identified to contain polymorphisms, allowing estimation of ASE. Genes were considered to show ASE when the number of associated reads assigned to B73 or PT transcripts was significantly different ( ^2^ test against an equal number of counts; p < 0.05; Bonferroni correction for multiple tests) and the absolute log2-transformed ratio of PT/B73 reads was > 1.

### Gene Ontology annotation, enrichment analyses and comparison of ASE genes to published data

Candidate ASE genes were assigned to Gene Ontology categories (release 52 available at ftp://ftp.gramene.org/pub/gramene). Obsolete annotations were replaced by the corresponding “consider” or “replaced_by” category(ies) in the ontology file (go.obo) available at http://www.geneontology.org/ (dated 2016-09-19). Categories associated with at least 10 genes were considered in further analysis. Enrichment analyses were performed comparing ASE candidates against the 9256 polymorphic gene set, using the Bingo (Maere, Heymans & Kuiper, 2005) Cytoscape (Shannon et al., 2003) plugin, controlling for multiple tests using Benjamini and Hochberg False Discovery Rate at 1%.

Candidate ASE genes were cross-referenced to a published study describing transcriptional responses in maize seedlings exposed to cold, heat, salt and UV stresses (Makarevitch et al., 2015). Although a number of inbred lines were analyzed in the Makarevitch study, only the B73 data was used in the comparison with the B73xPT transcriptome. Genes were considered to show a constitutive stress response (CR) with respect to a given stress when: 1) identified as ASE; 2) responding significantly to stress in the Makarevitch study (absolute log2 fold change >1; called as significant in the Makarevitch study; calls “up” or “on” in the published study were considered here as “up”, similarly, “down” or “off” were considered as “down”); 3) the sign of ASE was concordant with the sign of stress response.

Fst values for population level differentiation between Mesoamerican and South American highland and lowland maize populations (Takuno *et al*., 2015) were obtained from https://github.com/rossibarra/hilo_paper/tree/master/fst; where multiple SNPs are associated with a single gene, the values reported correspond to the SNP showing the highest Fst in Mesoamerica.

### Reconstruction of a gene co-expression network

Publicly available maize Affymetrix microarray data was downloaded from the ArrayExpress website (http://www.ebi.ac.uk/arrayexpress/; experiments E-GEOD-10023, E-GEOD-12770, E-GEOD-12892, E-GEOD-18846, E-GEOD-19785, E-GEOD-22479, E-GEOD-28479, E-GEOD-31188, E-GEOD-40052, E-GEOD-41956, E-GEOD-48406, E-GEOD-48536, E-GEOD-54310, E-GEOD-59533, E-GEOD-69659, E-MEXP-1222, E-MEXP-1464, E-MEXP-1465, E-MEXP-2364, E-MEXP-2366, EMEXP-2367, E-MEXP-3992). Low quality CEL files identified using the arrayQualityMetrics (Kauffmann, Gentleman & Huber, 2009) R package were discarded. Using the sample data relationship file (sdrf) associated with each experiment, samples for B73 leaves were selected, resulting in a high quality, homogeneous dataset of 165 CEL files.

Probeset sequences for the maize Affymetrix microarray were aligned using seqmap (Jiang & Wong, 2008) to the AGPv3.22 transcripts with no mismatches allowed, and probesets whose probe sequences did not align or aligned to transcripts corresponding to more than one locus were discarded. Probesets that were represented by less than 4 probe sequences were also discarded. This resulted in a list of 11299 probesets that unambiguously matched one locus. The list of 11299 probesets was used to create a custom chip definition file (CDF) using the ArrayInitiative python package (http://wellerlab.uncc.edu/ArrayInitiative/), and to filter the original Affymetrix Maize.probe_tab file to create a custom probe_tab file. The custom CDF and custom probe_tab file were then used to create the corresponding cdf and probe_tab R packages using the makecdfenv (Irizarry *et al*., 2006) and AnnotationForge (Carlson and Pages, 2017) R packages, respectively. The microarray name in the 165 CEL files was then modified to match the custom cdf and probe_tab packages name, and these modified CEL files were normalized using gcrma (Wu and Gentry, 2017). The resulting normalized dataset was then used as input for the ARACNE algorithm (Margolin et al., 2006a); (Margolin et al., 2006b), and inference was carried out for the 7 ASE and stress-responsive transcription factors (see Results) at DPI 0.1 as previously described (Chávez Montes et al., 2014). Enrichment analysis of the 1938 TF targets gene set was done as described above against the 11299 genes represented in the microarray.

## RESULTS

### A total of 2386 genes exhibited allele specific expression in the B73xPT F1 hybrid

To identify regulatory variation associated with stress-related genes, high throughput sequencing was used to quantify transcript abundance in leaves harvested from an F1 seedling generated from the cross between the Mexican highland landrace PT and the reference line B73. Alignment to the B73 reference gene models identified 9256 genes containing at least one sequence variant that could be used to distinguish the products of B73 and PT alleles. For 2386 (26%) of these 9256 polymorphic transcripts, the number of reads corresponding to the B73 allele differed significantly (p < 0.05; Bonferroni correction for multiple tests) from the number of reads corresponding to the PT allele with an absolute log2 fold change >1, and these genes were considered to exhibit allele specific expression (ASE; Supplementary Data 2 [F1_counts]). For 1412 (59%) of the ASE candidate genes, accumulation of the PT transcript was lower than that of the B73 transcript (log2 PT/B73 < -1; hereafter, “PT-down”), while for the remaining 974 (41%) of the ASE candidates, the PT transcript was accumulated at higher levels (log2 PT/B73 > 1; hereafter, “PT-up”).

To obtain an overview of the ASE candidates, a Gene Ontology (GO) analysis was performed. The set of 2386 ASE candidates was not enriched for any specific GO categories with respect to the 9256 polymorphic gene set, but, nonetheless, many individual genes belonged to biological processes categories related to stress responses, including responses to heat (GO: 0009408), cold (GO: 0009409) and salt (GO: 0009651) (Fig. 1). Overall, 52 biological process categories were represented by at least 10 genes. Of these, 38 (73%) were PT-down (based on the median log2 PT/B73 of the associated genes), and 11 (21%) were PT-up, and the remaining 3 categories had a median log2 PT/B73 close to 0 (Supplementary Data 3 [ASE_loci_GO_P]). A similar pattern was observed for molecular function categories: 57 categories were associated with at least ten ASE genes, 42 PT-down, 12 PT-up and 3 showing no trend (Supplementary Data 4 [ASE_loci_GO_F]).

**Figure 1.**
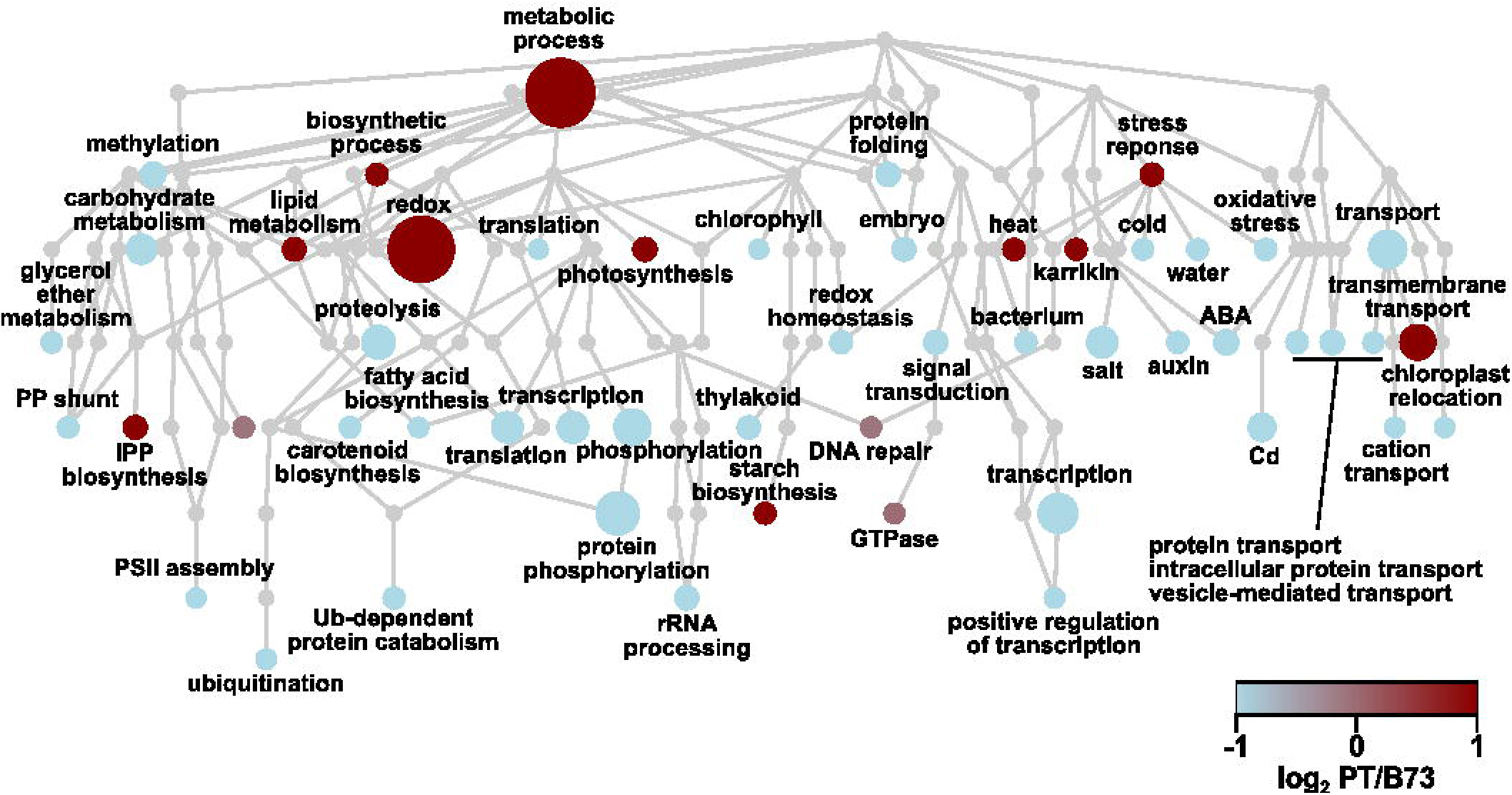
ASE candidate genes are assigned to a range of biological process Gene Ontology categories. Hierarchical tree of Gene Ontology biological process categories represented in ASE loci. Nodes represent categories, with the root GO:0008150 *biological process* as the uppermost node. Edges represent the parent-child (*i.e.* “is _a”) relationship between categories. Node color indicates the median ASE (log2 PT/B73) for the genes in the category, with light blue indicating negative values and dark red indicating positive values. Node size is proportional to the number of loci assigned to corresponding category. Some category names were abbreviated for clarity.**[2 COLUMNS]**

### A total of 277 genes showed constitutive stress responses

To identify evidence of a constitutive stress response (CR) in PT, the ASE gene set was compared with a previous study reporting changes in the transcriptome of B73 seedlings exposed to cold, heat, salt or UV treatments (Makarevitch *et al*., 2015). A total of 1407 stress responsive genes identified in the Makarevitch study were present also in the 9256 polymorphic gene set for which ASE had been evaluated (Supplementary Data 2 [F1_counts]). Of these 1407 genes, 432 (31%) showed ASE, a slight enrichment compared with the 2386 (26%) ASE genes in the 9256 polymorphic gene set as a whole ( ^2^ = 15.7, d.f. =1, p < 0.001). From this 432 gene set, a gene was considered to exhibit CR in PT if the sign of ASE was concordant with the sign of B73 stress response: *i.e.* PT-up and induced by stress in B73, or PT-down and repressed by stress in B73. On this basis, a set of 277 CR candidates was identified (Fig. 2A-D; Supplementary Data 5 [Maka_can_annot]). The majority of these 277 genes respond to two or more stress treatments (Fig.3A, C), but often in different directions such that most present stress-specific CR (Fig. 3B, C): 194 were identified as showing CR with respect to one treatment, 62 with respect to two, 17 with respect to three, and 4 with respect to all four (Fig. 3C). Of the 277 genes, 92 showed CR with respect to cold, 65 with respect to heat, 136 with respect to salt, and 92 with respect to UV (Fig. 3B). The number of CR genes with respect to any given stress was proportional to the number of genes responding to that stress in the 1407 polymorphic gene set (χ^2^ = 4.4, d.f. = 3, p = 0.22), and there was no indication of an enrichment for CR with respect to any one of the four treatments. In contrast to the complete ASE gene set, the majority of the 277 CR genes were PT-up (181 PT-up, 96 PT-down; Supplementary Data 5 [Maka_can_annot]), reflecting a bias also present in the 1407 gene set, although this general trend was not observed when the UV treatment was considered alone, where the majority of CR genes were PT-down (Fig. 2D).

**Figure 2.**
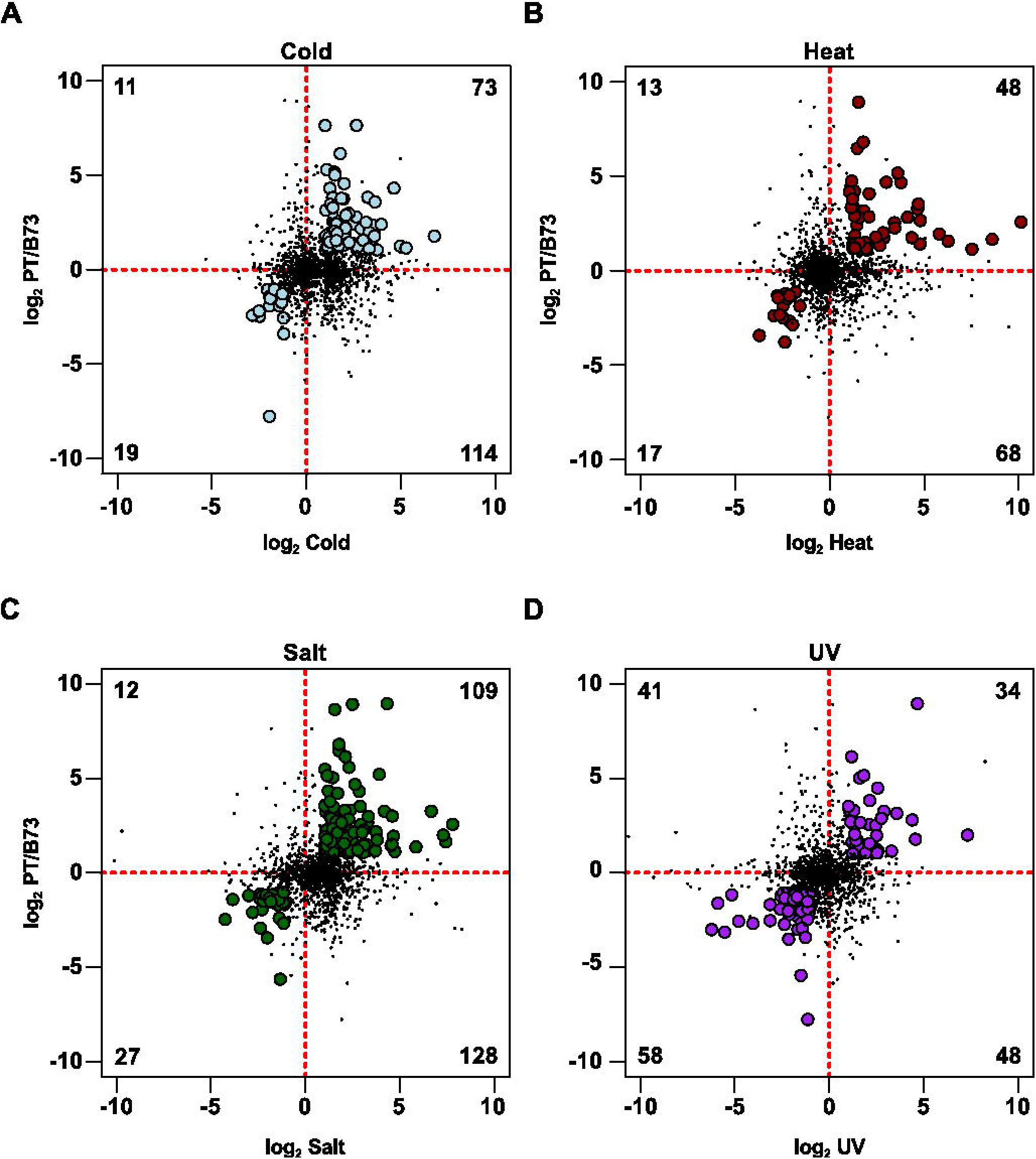
ASE identifies constitutive stress response in PT with respect to B73. ASE (log2 PT/B73) in control F1 leaves for the 1407 sequence variant, stress-responsive gene set against B73 stress response (log2 stress/control) for A) cold, B) heat, C) salt and D) UV treatments as reported in the Makarevitch dataset. Numbers in each quadrant represent the count of genes called as significant in ASE and stress comparisons. In each plot, the quadrants represent (clockwise from upper left) genes up ASE / down stress, up ASE / up stress, down ASE / up stress, down ASE / down stress. Genes called as up ASE / up stress or down ASE / down stress are considered canalized and are shown as filled circles. Other genes are shown as points. Axes through the origin are shown as red dashed lines. A number of genes outside the axis range are not shown, but are considered in the gene count. **[TWO COLUMNS]**

**Figure 3.**
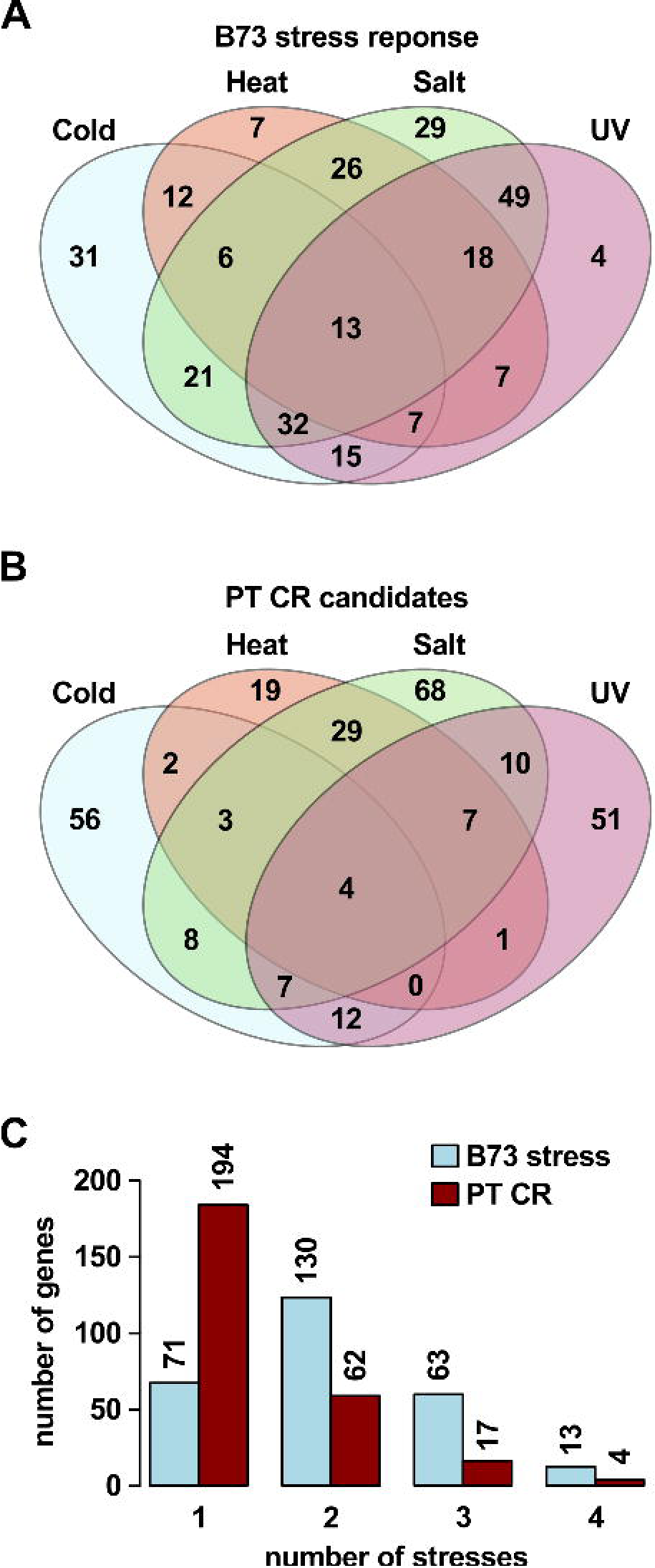
CR candidates may respond to multiple stresses in B73. **A)** Number of genes from the 277 CR gene set that responded to cold, heat, salt, UV or a combination of stresses in the Makarevitch B73 study. **B)** Number of genes called as CR in PT with respect to each stress from the same 277 gene set. **C)** Counts with respect to number of stresses of genes in A and B. Numbers above bars give counts. **[ONE COLUMN]**

### Hormone related genes and transcription factors showed constitutive stress responses in PT

A primary aim of the analysis was the definition of a small number of candidate genes for future functional analysis. For this purpose, the CR candidate genes were cross-referenced with the classical maize gene list, a curated set of 4908 well-annotated genes, many linked with existing functional data (www.maizegdb.org/gene_center/gene). Of the 277 CR candidate genes, 48 were present in the classical gene list (Fig. 4; Supplementary Data 5 [Maka_can_annot]), including 9 genes associated with hormone homeostasis (Table 1) and 12 transcription factors (TFs; Table 2; (Jin et al., 2017)) that were considered of special interest. The 277 CR candidates were cross referenced with a published study of population level differentiation between Mesoamerican and South American highland and lowland maize (Takuno *et al.*, 2015). Twenty-two of the 277 CR candidates showed significant Fst (p < 0.01) between highland and lowland Mesoamerican populations, including the hormone associated gene *Czog1* (GRMZM2G168474; Supplementary Data 5 [Maka_can_annot]). To gain insight into potential TF targets and their role in stress responses, a gene co-expression network for the CR TFs was generated using available maize Affymetrix microarray data and the ARACNE algorithm. Seven of the 12 TFs were unambiguously identified in the maize Affymetrix microarray probeset, and were co-expressed with 1938 genes (Supplementary Data 6 [tfs_ASE_01_suppl]). Co-expressed genes represent potential targets of TF action, and, as such, may not themselves exhibit ASE. Indeed, of the 1938 genes associated with the 7 TFs, 1097 were present in the polymorphic gene set, but only 239 showed ASE. A total of 344 of the 1938 co-expressed genes (17%) were responsive to one or more stress treatments in the Makarevitch dataset (Fig. 5). A GO analysis detected enrichment in the 1938 gene co-expression set with respect to translation, photosynthesis and non-mevalonate isoprenoid pathway categories (Supplementary Data 7 [Bingo_aracne]).

**Figure 4.**
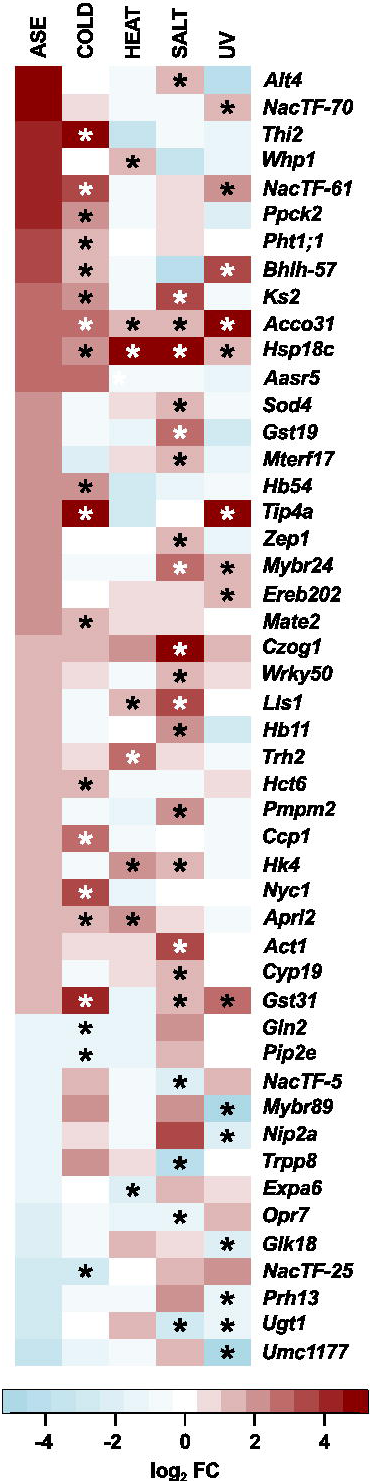
Classical CR candidate genes. Heatmap representation of ASE (log2 PT/B73) and B73 response to cold, heat, salt and UV stress (log2 stress/control) as reported in the Makarevitch dataset. Asterisks (*) in the stress columns indicate a given gene was called as CR with respect to that stress. **[ONE COLUMN]**

**Table 1.** ASE and stress-responsive hormone-related genes. List of genes involved in hormone biosynthesis, transport or catabolism present in the 277 CR gene set. ASE call indicates biased expression of the PT allele (1) or B73 allele (-1). Response to stress indicates the name of the stress for which the gene was called as differentially expressed in the Makarevitch dataset. Constitutive response indicates the stress condition for which the sign of the ASE call and the stress response coincide.

| Gene id | Symbol | Molecular function | Hormone | ASE call | Response to stress | Constitutive response |
| --- | --- | --- | --- | --- | --- | --- |
| GRMZM2G070563 | --- | auxin efflux carrier | auxin transport | 1 | heat, salt, uv | heat, salt |
| GRMZM2G072632 | --- | auxin efflux carrier | auxin transport | 1 | heat, salt, uv | heat, salt |
| GRMZM2G112598 | --- | auxin efflux carrier | auxin transport | 1 | heat, salt, uv | heat, salt |
| GRMZM2G475148 | --- | auxin efflux carrier | auxin transport | 1 | heat, salt | heat, salt |
| GRMZM2G072529 | *Acco31* | 1-aminocyclopropane-1-carboxylate oxidase | ethylene biosynthesis | 1 | cold, heat, salt, uv | cold, heat, salt, uv |
| GRMZM2G020761 | -- | putative cytochrome P450 (castasterone C-26 hydroxylase) | brassinosteroid catabolism | -1 | cold, salt, uv | cold, uv |
| GRMZM2G148281 | *Opr7* | 12-oxo-phytodienoic acid reductase | jasmonate biosynthesis | -1 | salt, uv | salt |
| GRMZM2G168474 | *Czog1* | *cis*-zeatin O-glucosyl transferase | cytokinin<br>homeostasis | 1 | salt | salt |

**Table 2.**
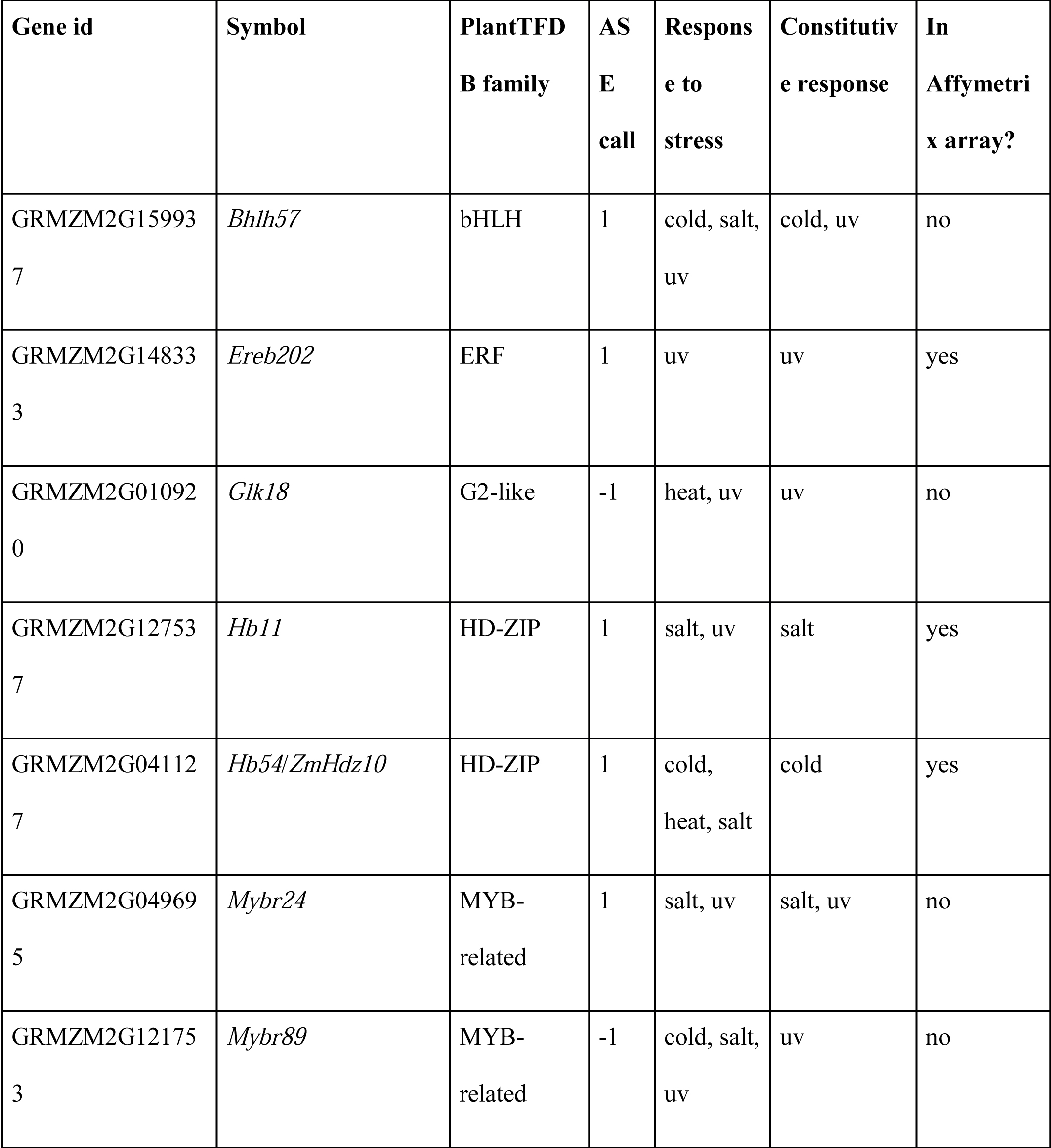
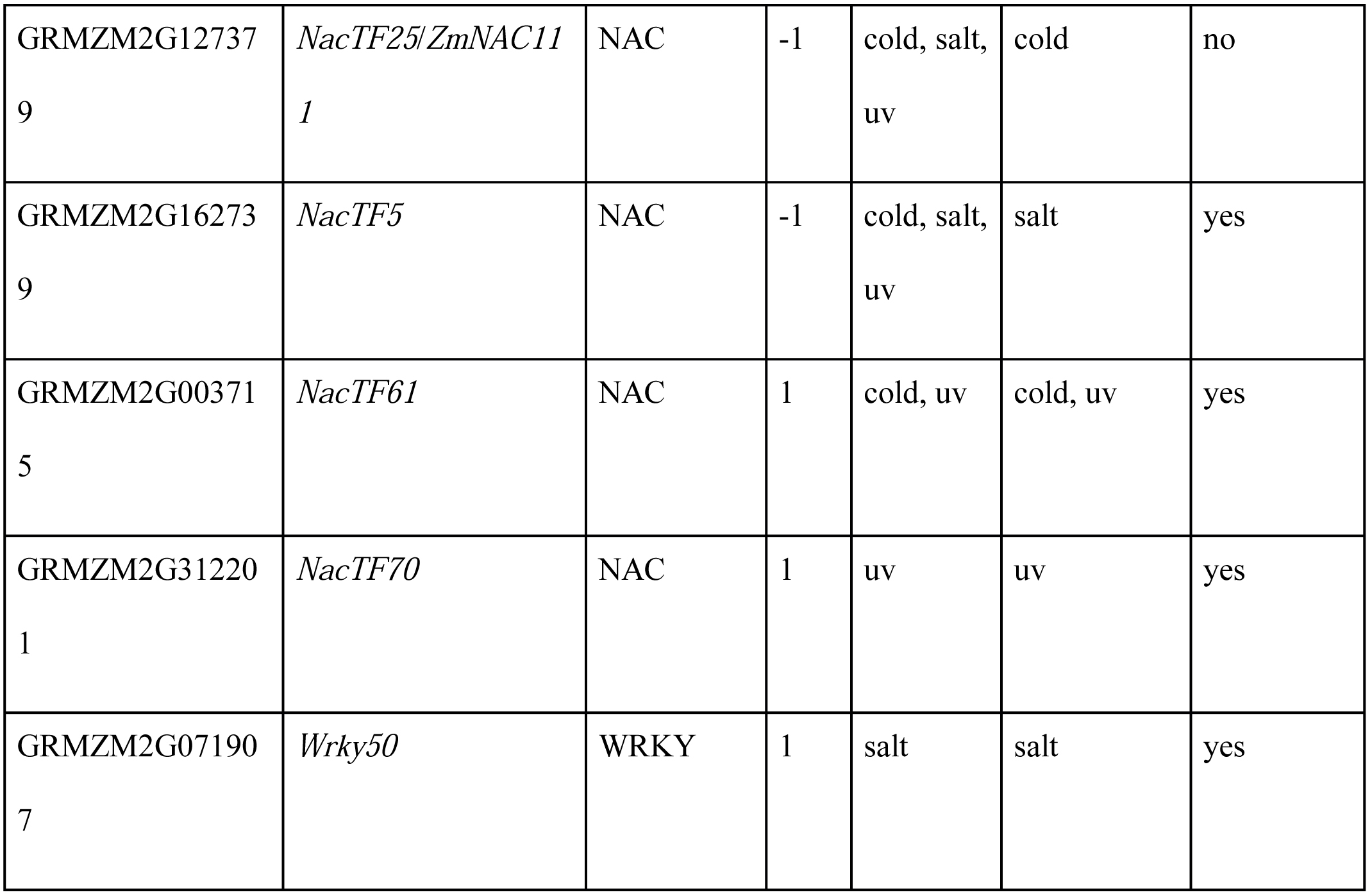
ASE and stress-responsive TFs. List of TFs present in the 277 CR gene set. PlantTFDB family indicates the TF family according to the PlantTFDB (Jin et al. 2017). ASE call indicates biased expression of the PT allele (1) or B73 allele (-1). Response to stress indicates the name of the stress for which the gene was called as differentially expressed in the Makarevitch dataset. Constitutive response indicates the stress condition for which the sign of the ASE call and the stress response coincide. In Affymetrix array indicates of the TF is represented in the maize Affymetrix microarray.

| Gene id | Symbol | PlantTFDB family | ASE call | Response to stress | Constitutive response | In Affymetrix array? |
| --- | --- | --- | --- | --- | --- | --- |
| GRMZM2G159937 | *Bhlh57* | bHLH | 1 | cold, salt, uv | cold, uv | no |
| GRMZM2G148333 | *Ereb202* | ERF | 1 | uv | uv | yes |
| GRMZM2G010920 | *Glk18* | G2-like | -1 | heat, uv | uv | no |
| GRMZM2G127537 | *Hb11* | HD-ZIP | 1 | salt, uv | salt | yes |
| GRMZM2G041127 | *Hb54/ZmHdz10* | HD-ZIP | 1 | cold, heat, salt | cold | yes |
| GRMZM2G049695 | *Mybr24* | MYB-related | 1 | salt, uv | salt, uv | no |
| GRMZM2G121753 | *Mybr89* | MYB-related | -1 | cold, salt, uv | uv | no |
| GRMZM2G12737<br>9 | *NacTF25/ZmNAC11*<br>*1* | NAC | -1 | cold, salt,<br>uv | cold | no |
| GRMZM2G16273<br>9 | *NacTF5* | NAC | -1 | cold, salt,<br>uv | salt | yes |
| GRMZM2G00371<br>5 | *NacTF61* | NAC | 1 | cold, uv | cold, uv | yes |
| GRMZM2G31220<br>1 | *NacTF70* | NAC | 1 | uv | uv | yes |
| GRMZM2G07190<br>7 | *Wrky50* | WRKY | 1 | salt | salt | yes |

**Figure 5.**
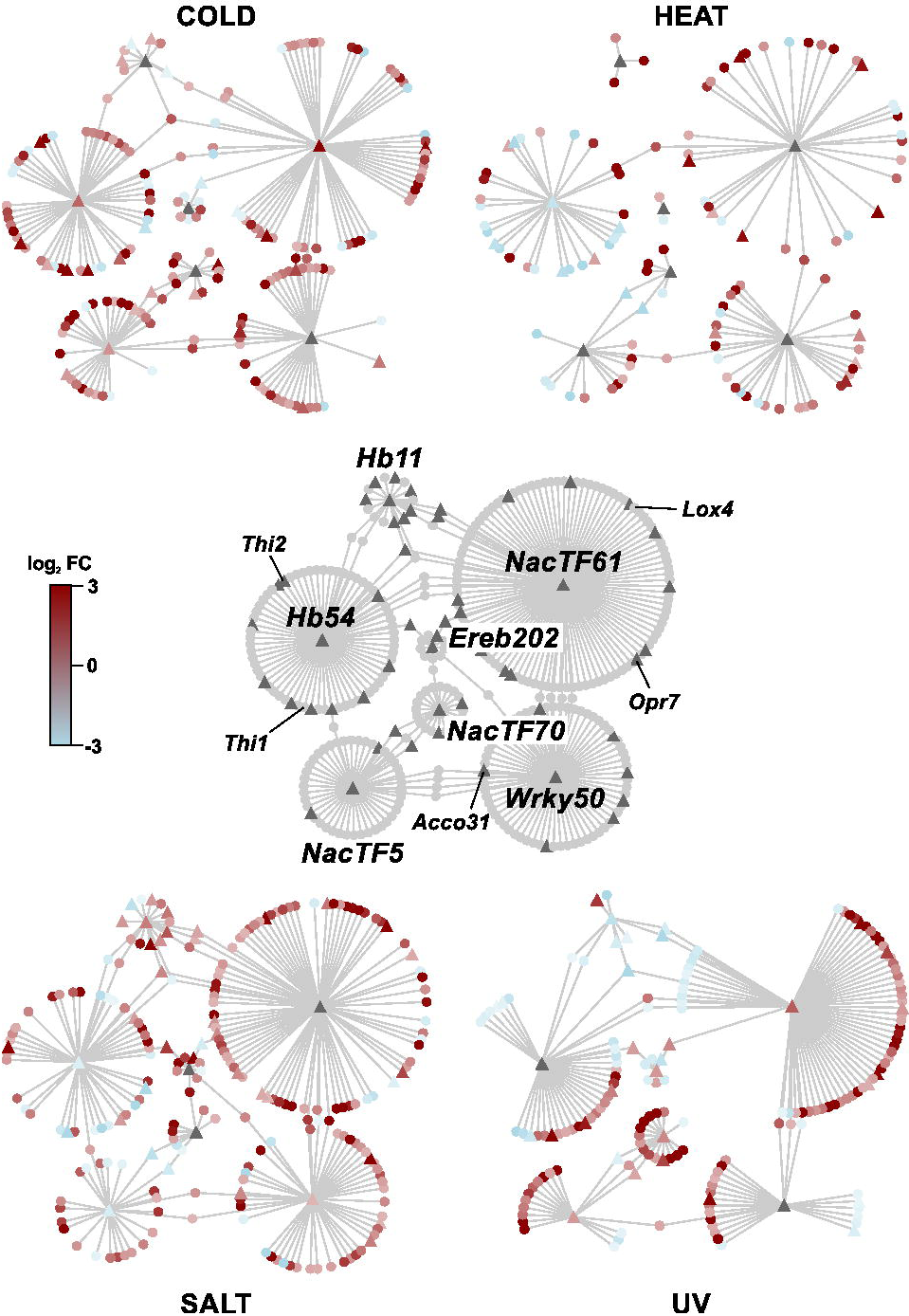
Co-expression networks for CR TFs and their putative stress-responsive targets. Nodes represent genes and edges represent co-expression as calculated by the ARACNE algorithm at DPI 0.1. The center panel presents a network of seven CR TFs (labeled centres of circles) with their co-expressed, stress-responsive (genes called up/on or down/off in the Makarevitch dataset) putative targets. Triangles indicate genes that were called as presenting ASE. The network was filtered to retain only co-expressed genes responsive cold, salt, UV or heat treatments. In the filtered networks the red and blue colors indicate up or down regulation (as log2 FC from the Makarevitch dataset), respectively, under the corresponding stress.**[TWO COLUMNS]**

## DISCUSSION

From a starting set of 9256 polymorphic genes, we identified 2386 genes presenting allele specific expression (ASE) in seedling leaves of an F_1_ B73xPT hybrid individual. Comparison of our ASE gene list with a published dataset reporting B73 stress responses (Makarevitch et al., 2015) identified a subset of 277 (out of 432) constitutive stress response (CR) candidate genes exhibiting a bias in transcript accumulation between PT and B73 alleles that mirrored the B73 response to one or more stress treatments. We did not observe an enrichment in GO term assignments in either our ASE gene set or our CR gene set. Nonetheless, given that ASE is assaying *cis*-acting variation, a small number of genes associated with a given GO term may have biological significance. The ASE gene set showed a bias towards lower expression of the PT allele, reflected in the observation that the median value of ASE for the majority of GO categories associated with ASE genes was also negative. Contrary to this trend, the subset of 277 selected CR candidates showed a bias towards higher expression of the PT allele (181 of 277 presented higher expression of the PT allele), also reflected in the 1407 polymorphic genes that overlapped with the Makarevitch set.

The bulk of the CR gene set (206 of 277) responded to two or more stresses in the Makarevitch B73 data, although in the majority (194 of 277) of cases the CR itself was with respect to a single stress only (Fig. 3), indicating that in many cases the sign (up/down) of the response in B73 differed between stresses (Supplementary Data 5 [Maka_can_annot]). In total, of the 1407 polymorphic and stress-responsive genes in the working set, 511 genes induced in B73 under at least one stress treatment were repressed under at least one other (Supplementary Data 2 [F1_counts]). By definition, a gene could not show CR with respect to both of two different stresses if the B73 responses were opposing. There was no evidence that genes showing opposing stress responses in B73 were less likely to show CR in PT -- indeed, such genes were actually better represented in the 277 CR gene set (56%) than in the 1407 polymorphic and stress-responsive gene set (36%). As such, many ASE events may appear contradictory with respect to any given stress, *i.e.* PT-up ASE in genes repressed by B73 under stress, or PT-down ASE in genes induced by B73, especially in the context of cold and UV treatments, against which PT is considered to be well adapted. The spatio-temporal dynamics of stress responses, however, are complex (*e.g.* (Secco et al., 2013), and the resolution of the present analysis, based on single time points and tissues, is limited. For example, the previously characterized salt associated HD-ZIP transcription factor *Hb54* (also named *ZmHdz10*, GRMZM2G041127; (Zhao et al., 2011); (Zhao et al., 2014) showed PT-up ASE, but was repressed by salt treatment in the Makarevitch dataset, and consequently not considered to show CR. In this case, however, an additional functional study reports *Hb54* to indeed be induced by salt treatment (Zhao et al., 2014), albeit at a different time point, and with a different treatment than that applied in the Makarevitch study (300mM NaCl for 20hrs in Makarevitch *et al.*; 200mM NaCl for 3-12hrs in Zhao *et al*.). The study of Zhao and colleagues reports also that constitutive expression of *Hb54* in *Arabidopsis* and rice increases ABA sensitivity and tolerance to drought and salt stress. In light of these data, PT-up ASE of *Hb54* may indeed have biological relevance, reflected by the number and nature of associated co-expression candidates (Fig. 5). In the absence of further characterization, it would be premature to discount the potential phenotypic impact, or adaptive value, of other examples where ASE in PT is opposed to the B73 stress response reported in the Makarevitch data.

Previous studies have highlighted the importance of *cis*-acting regulatory variation in driving diversity in plant stress responses (*e.g.* (Waters et al., 2017). The generation of novel physiological strategies to confront stress conditions may be most efficient when a change in the regulation of a single gene has multiple, coordinated downstream consequences. Mechanistically, two functional categories of clear interest are hormones, systemic regulators of physiology at the whole plant level, and transcription factors (TFs), with their capacity to impact multiple downstream targets through a regulatory cascade. The 277 CR gene list includes eight hormone-related genes (Table 1), including genes implicated in the metabolism of cytokinin (*Czog1* ; (Martin et al., 2001), jasmonate (*ZmOpr7*; (Yan et al., 2012) and ethylene (*Acco31*; (Gallie & Young, 2004); (Avila et al., 2016). Additional CR candidates included *Ks2* (GRMZM2G093526; *ZmKSL5*), a gene related to the *ent*-kaurene synthase required for gibberellin biosynthesis, but more likely involved in the more specialized kauralexin A series biosynthesis pathway (Fu et al., 2016), and *Thi2* (GRMZM2G074097), encoding a thiamine thiazole synthase activity required for synthesis of the thiazole moiety during the production of thiamin (vitamin B1; (Woodward et al., 2010). With regard to the latter candidate, B vitamins, although not strictly plant hormones, can play an analogous role in whole plant physiology in the face of stress (Hanson et al., 2016). Thiamin application has been reported to alleviate the impact of abiotic stress in a number of crops, including maize (*e.g.* (Kaya et al., 2015), and thiamin synthesis has been proposed as a target for transgenic biofortification (*e.g.* (Dong, Stockwell & Goyer, 2015). Identification of PT-up ASE associated with *Thi2* represents a compelling target for further analysis. Interestingly, both *Thi2* and the related gene *Thi1* (GRMZM2G018375) were also co-expressed with the PT-up ASE drought and salt associated HD-ZIP TF *Hb54* ((Zhao et al., 2014); Table 2; Fig. 5; Supplementary Data 6). Interestingly, the CR candidates *Czog1* and *Ks2* were reported previously to show significant population level differentiation between highland and lowland mesoamerican maize populations (F_st_; p = 0.004, p = 0.04, respectively; (Takuno et al., 2015), indicating that variation at these loci may indeed play a role in local adaptation.

In total, twelve TFs were present in the 277 CR candidate gene set (Table 1), including four NAC TFs. The NAC TFs are a plant-specific family implicated broadly in abiotic stress responses (Nakashima et al., 2012); (Puranik et al., 2012); (Nuruzzaman, Sharoni & Kikuchi, 2013); (Nakashima, Yamaguchi-Shinozaki & Shinozaki, 2014), previously proposed as a target for engineering multiple stress tolerance (Shao, Wang & Tang, 2015). The potential role of NAC TFs in a generalized stress response is reflected by the observation that the candidates *ZmNacTF5* (GRMZM2G162739)*, ZmNacTF25* (also named *ZmNac111*, GRMZM2G127379; (Mao et al., 2015)*, ZmNacTF61* (GRMZM2G003715) and *ZmNacTF70* (GRMZM2G312201) responded to 3, 3, 2 and 1 stress treatments, respectively (Table 2). The genes *ZmNacTF5* and *ZmNacTF25* showed PT-down ASE, and CR with respect to salt and cold, respectively, while the genes *ZmNacTF61* and *ZmNacTF70* showed PT-up ASE and CR with respect to cold and UV, respectively. In B73, insertion of a miniature inverted-repeat transposable element (MITE) in the *ZmNacTF25* promoter has been reported previously to be associated with reduced gene expression (relative to a number of tropical lines) and increased susceptibility to drought (Mao et al., 2015). The accumulation of *ZmNacTF25* transcripts in B73, however, is reduced under cold in the Makarevitch dataset, indicating a potential trade-off between temperate and tropical lines, and possible relevance of the PT-down ASE in the highland niche. The gene *ZmNacTF61* was notable for strong PT-up ASE (log2 PT/B73 = 3.26 and 2.15) , up-regulation under both cold and UV stress, and association with a large number (116) of strongly cold- and UV-induced co-expression candidates, including the jasmonate biosynthetic genes *Opr7* and *Lox4* (Fig. 4; Fig. 5; Supplementary Data 6).

Candidate CR genes presented here were identified on the basis of ASE under benign conditions. Investigation of the degree to which ASE is maintained under stress conditions is required to determine whether expression of these candidates has indeed been canalized to a constitutively responsive state, or whether expression of the PT alleles remains plastic, albeit with an expression level different from B73. Nonetheless, the potential to identify relevant *cis*-regulatory variation through exploration of the transcriptome under benign conditions presents an attractive avenue to investigate stress response and local adaptation. A number of the candidates identified here suggest testable predictions regarding hormone accumulation and expression of candidate TF targets in the PT landrace. In a number of cases, ASE was observed in genes reported previously to show significant genetic differentiation between lowland and highland Mexican maize populations, offering further evidence of a link to adaptation to the highland niche (Supplementary Data 5 [Maka_can_annot]). A recent study in monkey flower (*Mimulus guttatus*) using ASE analysis to compare locally adapted coastal and inland accessions has found *cis*-regulatory effects to be the main driver for regulatory variation, providing a precedent for the approach proposed here (Gould, Chen & Lowry, 2017). Validation of specific candidate genes will require functional characterization, but it is anticipated that this will be greatly facilitated by continued development of resources for maize reverse genetics and the forthcoming availability of introgression lines derived from Mexican highland maize (Matt Hufford and Ruairidh Sawers, unpublished data).

## CONCLUSIONS

Expression differences were observed between PT and B73 alleles under benign conditions that mirror the B73 response to cold, heat, salt and/or UV treatments. The observed patterns of expression indicate the presence of *cis*-acting regulatory variation differentiating the PT landrace from the B73 reference inbred. Regulatory variants linked to classical genes associated with signaling and stress-responses potentially contribute to the adaptation of PT to the Mexican highland environment.

## ACKNOWLEDGEMENTS

We acknowledge Patrice Dubois for assistance in the generation of F1 seed stock, and Patrick Schnable and Cheng Ting Yeh for generation of transcriptome data.

